# Efficiency enhancement of microparticles seeding density on the inner surface of polymer hollow microfibers using microfluidics

**DOI:** 10.1101/2023.01.16.524327

**Authors:** Saurabh S. Aykar, Nicole N. Hashemi

## Abstract

Lateral displacement of microparticles suspended in a viscoelastic fluid flowing through a microfluidic channel occurs due to an imbalance in the first (N1) and second (N2) normal stress differences. Here, we studied the lateral displacement of fluorescent microparticles suspended in a polyethylene glycol (PEG) solution in a two-phase flow with aqueous sodium alginate, flowing through a unique microfluidic device that manufactures microparticles seeded alginate-based hollow microfibers. Parameters such as concentration of the aqueous sodium alginate and flow rate ratios were optimized to enhance microparticle seeding density and minimize their loss to the collection bath. 4 % w/v aqueous sodium alginate was observed to confine the suspended microparticles within the hollow region of microfibers as compared to 2 % w/v. Moreover, the higher flow rate ratio of the core fluid, 250 *μL min*^−1^ resulted in about 192 % increase in the microparticle seeding density as compared to its lower flow rate of 100 *μL min*^−1^. The shear thinning index (m) was measured to be 0.91 for 2 % w/v and 0.75 for 4 % w/v sodium alginate solutions. These results help gain insights into understanding the microparticle displacement within a viscoelastic polymer solution flowing through a microfluidic channel and motivate further studies to investigate the cellular response with the optimized parameters.

## Introduction

Lateral displacement of particles suspended in a fluid flowing through a microfluidic channel can be utilized in several applications such as cell sorting ^1–5^, cell encapsulation ^6–14^, sample preparation ^15–17^, etc. The particle displacement in a microfluidic channel could be achieved by imposing active or utilizing passive forces depending on the type of associated particles and their corresponding applications. The active methods include the external forces generated electrically ^18, 19^, optically ^20, 21^, and acoustically ^22–24^, while the passive methods utilize the forces generated from the fluid type, flow attributes, and microfluidic channel geometry. For applications involving particles that are sensitive to external forces, high throughput methods harnessing the invasive intrinsic forces for inducing lateral particle displacement are required. Since the passive methods do not require any external forces, they can be applied to applications that require a label-free and invasive approach.

Passive microfluidic methods, along with utilizing the forces derived from fluid flow attributes and microchannel geometry ^25, 26^, also depend on the physical properties of the suspended particles, such as size and rigidity ^27, 28^. For example, the lateral displacement of these particles within a microfluidic channel flow is directly proportional to their hydraulic diameter. The type of fluid flowing within the microchannel and the microchannel geometry and its cross-section govern the direction of the particle displacement. The non-Newtonian fluids give rise to additional forces such as first and second normal stress differences and elastic forces along with shear thinning or shear thickening effects. Unlike Newtonian fluids, the viscosities of these solutions aren’t constant and vary with the rate of shear strain. All these aforementioned forces resulting from the non-Newtonian fluid flow affect the particle displacement within the microchannel.

Cell or drug encapsulation is one of the applications where cells or drugs are targeted to specific regions of a hydrogel for several applications, such as controlled drug delivery ^29, 30^ or mimicking a specific architecture of a tissue or an organ. To drive these particles to the desired locations passively in the suspended fluid flowing within the microchannel, the forces acting on these particles should be accounted for. Here, the focus of this study is to investigate the particle displacement within a polymer solution flowing adjacent to another polymer solution in a straight rectangular microchannel. The results of this study could essentially be used to determine the desired cell seeding or encapsulation locations within a polymer hollow microfiber.

One of the applications is the microfluidic seeding of cells on the inner surface of polymer-based hollow microfibers, the process of which is expounded in this study using microparticles as representatives of cells. This cell seeding on the inner surface of hollow polymer microfibers using microfluidics has enabled us to mimic the structure of various microvascular systems in vitro. ^31,32^ The desired cells are suspended in the cytocompatible polymer solutions, either natural or synthetic, and a concentric fluid regime is achieved using microfluidic devices. The cell-suspended polymer solution eventually ends up in the core of the concentric fluid regime, and upon polymerization, cells get seeded on the inner surface of the polymer microfibers while the solution runs into the collection bath. In order to increase the final cell seeding density on the inner surface of hollow polymer microfibers, the cells must displace towards the interface of the core and the annulus solutions. In this study, different concentrations of polymer solutions with different flow rate ratios were investigated to increase the particle seeding density within the hollow polymer microfibers. The results of the study could be translated to increase the cell seeding density within the hollow polymer microfibers that could be further used to mimic various microvascular systems in-vitro.

## Background

Polymer fluids are considered viscoelastic fluids as they exhibit both viscous and elastic properties. These fluids exhibit features different from the Newtonian fluids, such as the Weissenberg climbing effect and the Barus effect. These properties are attributed to their viscous and elastic components. For instance, the viscosity of the viscoelastic fluids decreases with an increase in shear rate, unlike Newtonian fluids, where it only varies with temperature. This can be attributed to the fluid’s macromolecular behavior.

Particle migration across the streamlines in a viscoelastic flow largely depends on the first (N1) and second (N2) stress differences. N1 is a difference between two independent normal stresses along the flow and transverse to the flow. N1 is attributed to the material property *ψ*1, called as first normal stress difference coefficient, and is defined as 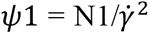, where 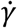 is rate of shear strain. N2 is defined as a difference between two independent normal stresses in the transverse direction of the flow, which gives rise to the material property *ψ*2, where 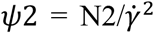. ^33–35^ Generally, N2 is an order of magnitude smaller than N1; however, in rectangular channels, the former induces circulation perpendicular to the flow direction. The elastic component of the viscoelastic fluid is governed by fluid relaxation time (*λ*) and is defined as the time require for the deformed fluid particles to restore to their equilibrium state. The fluid viscosity (*μ*) and relaxation time (*λ*), along with normal stress difference coefficients (*ψ*1 & *ψ*2) are used for the characterization of the viscoelastic flows.

In Poiseuille flow, the elastic force (Fel) derived from N1 drives the suspended particle towards the channel centerline, where the rate of shear strain is lowest due to the least velocity gradient present. This force largely depends on the particle size suspended in the flow and is directly proportional to it. Conversely, the fluid shear thinning effect drives the particles towards the wall where the high rate of shear strain exists. This dual phenomenon suggests that two equilibrium streamlines exist, which are governed by N1 and the shear thinning effect exhibited by the fluid. If the elastic component of the viscoelastic fluid is higher compared to the shear thinning, the suspended particles will travel towards the centerline, whereas if the latter is higher, they will migrate towards the walls of the channel. The dimensions of the microfluidic channel also play a crucial role in the migration of the particles. A dimensionless parameter called Blockage ratio (*β*) is defined as the ratio of characteristic lengths of particle and channel. At higher *β*, the particles tend to migrate towards the nearest walls of the channel. The particle velocity across the streamlines during migration to the equilibrium location depends on the fluid properties and the particle size.

## Results and Discussion

The fluorescent microparticles suspended in the core solution containing PEG were seeded on the inner surface of the alginate hollow microfibers using the previously developed microfluidic device ^32, 36^. In this study, the seeding density of the microparticles on the inner surface of the alginate hollow microfibers was enhanced by analyzing the concentration of the precursor solution, and flow rate ratios of these solutions. Conversely, the study focuses on minimizing the loss of these microparticles to the collection bath containing the polymerizing solution, which is critical when cells or biomolecules are used for seeding and are present in limited quantity. The microfluidic manufacturing process of seeding microparticles on the inner surface of the alginate hollow microfibers is illustrated in **Figure 1**.

**Figure 1.**
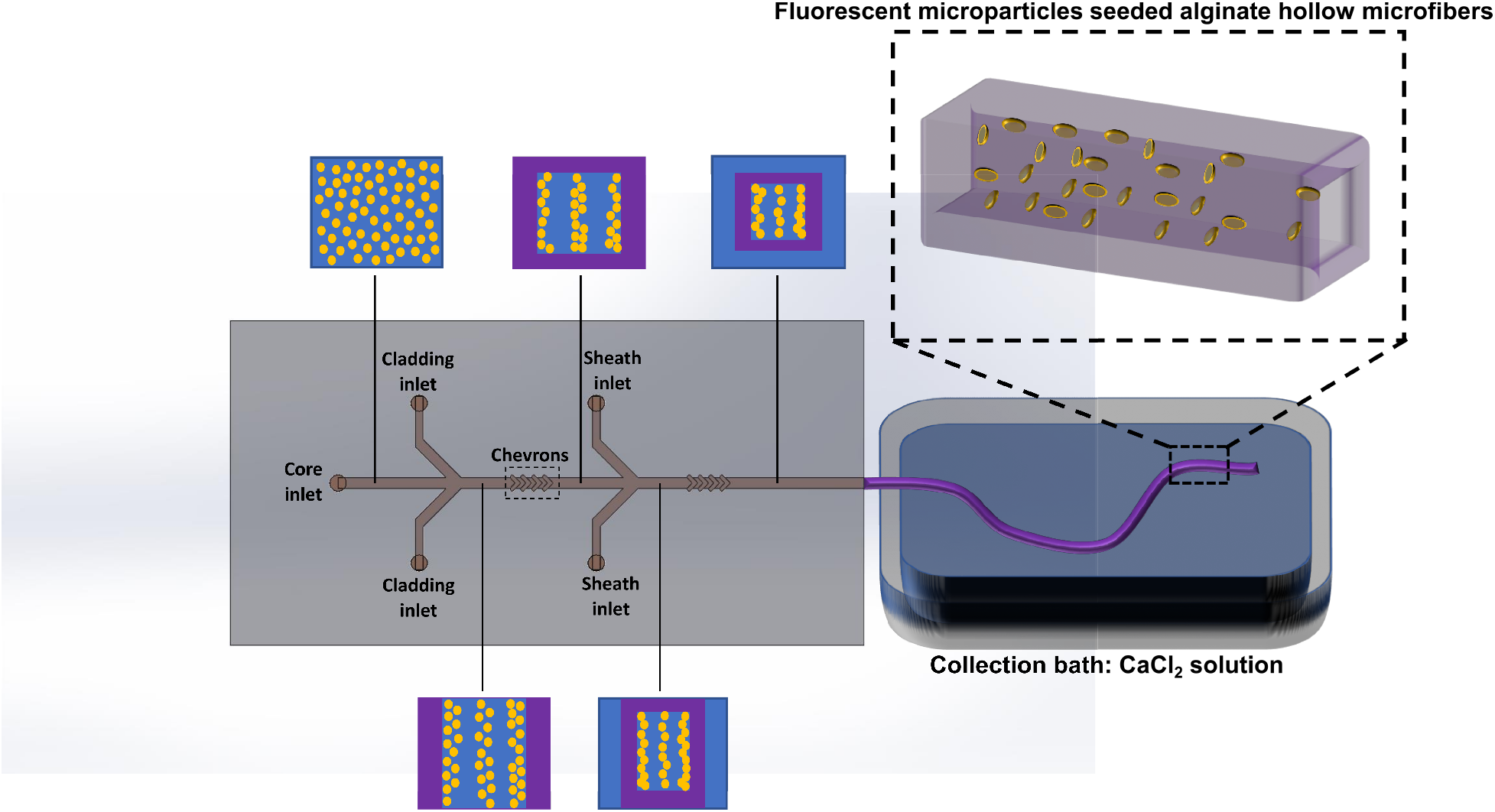
Schematic illustration of microparticles seeding on the inner surface of alginate hollow microfibers. The schematic cross-sectional view of the concentric fluid regime developed at distinct points along the central microchannel is shown. Blue and purple represent the PEG and alginate solutions, respectively, while the yellow dots represent the fluorescent microparticles. On the top right corner, the fluorescent particles are shown seeded on the inner surface of the alginate hollow microfibers.

### Effect of different concentrations of alginate on particle displacement

Initially, 2 % w/v alginate solution was used as a precursor for the seeding of microparticles that were suspended in the core solution containing PEG on the inner surface of the alginate hollow microfibers. It was observed that along with microparticles seeded on the inner surface of the alginate hollow microfibers, they were also encapsulated in the walls. This is evident from **Figure 2**, which represents the fluorescence images of the microparticles seeded with 2 % w/v alginate solution as a precursor. The fluorescent microparticles can be traced all along the width of the hollow microfibers. The encapsulation of the microparticles occurred for all the flow rate ratios. This showed that the 2 % w/v alginate solution could not confine the microparticles at the interface of the core and cladding solution. This could be attributed to the viscosity of the alginate solution being below a threshold value, that in turn was unable to confine the microparticles within the core fluid regime. To measure the reduction in the viscosity of the alginate solution due to shear thinning while flowing through the microchannel, the viscosities of 2 % w/v and 4 % w/v alginate solutions were measured over a range of velocities and are shown in **Figure 3**. The shear thinning index (m) for 2 % w/v and 4 % w/v alginate solutions was calculated to be 0.91 and 0.75, respectively. This shows that there was not any drastic changes in the viscosity observed during its flow within the microfluidic device. The average viscosity measured for 2 % w/v and 4 % w/v sodium alginate was 89.44 cP and 625.7 cP, respectively. This shows that the viscosity of 4 % w/v sodium alginate solution was considerably higher than its counterpart and did not change drastically during the flow within the microchannel. To confine the lateral displacement of microparticles within the boundaries of the concentric fluid regime containing the core solution, the viscosity of the alginate solution was increased by increasing its concentration.

**Figure 2.**
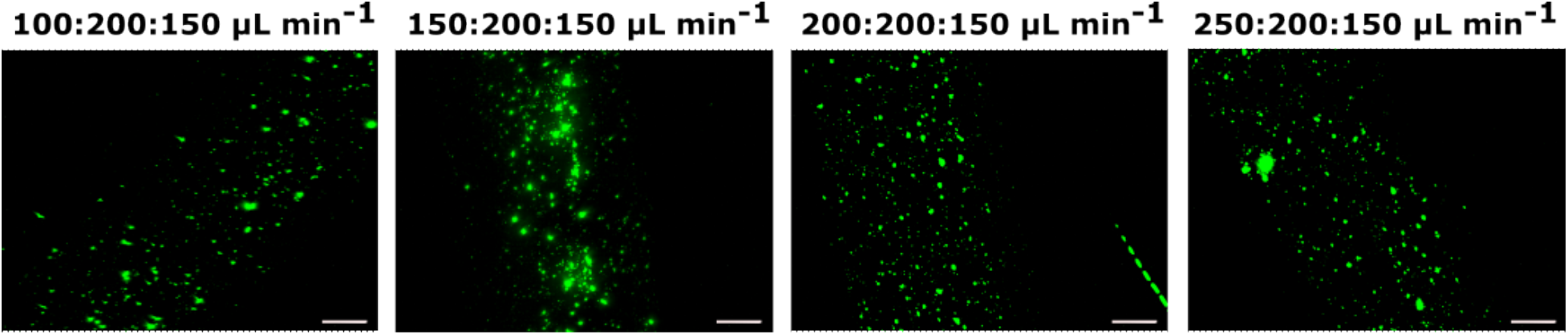
Fluorescence images of the microparticles seeded on the inner surface of alginate microfibers. 2 % w/v alginate solution was used as a precursor solution. The fluorescent microparticles were observed across the entire width of the alginate hollow microfibers for all flow rate ratios indicating their leakage into the walls of the microfibers. The scale bars are 100 μm.

**Figure 3.**
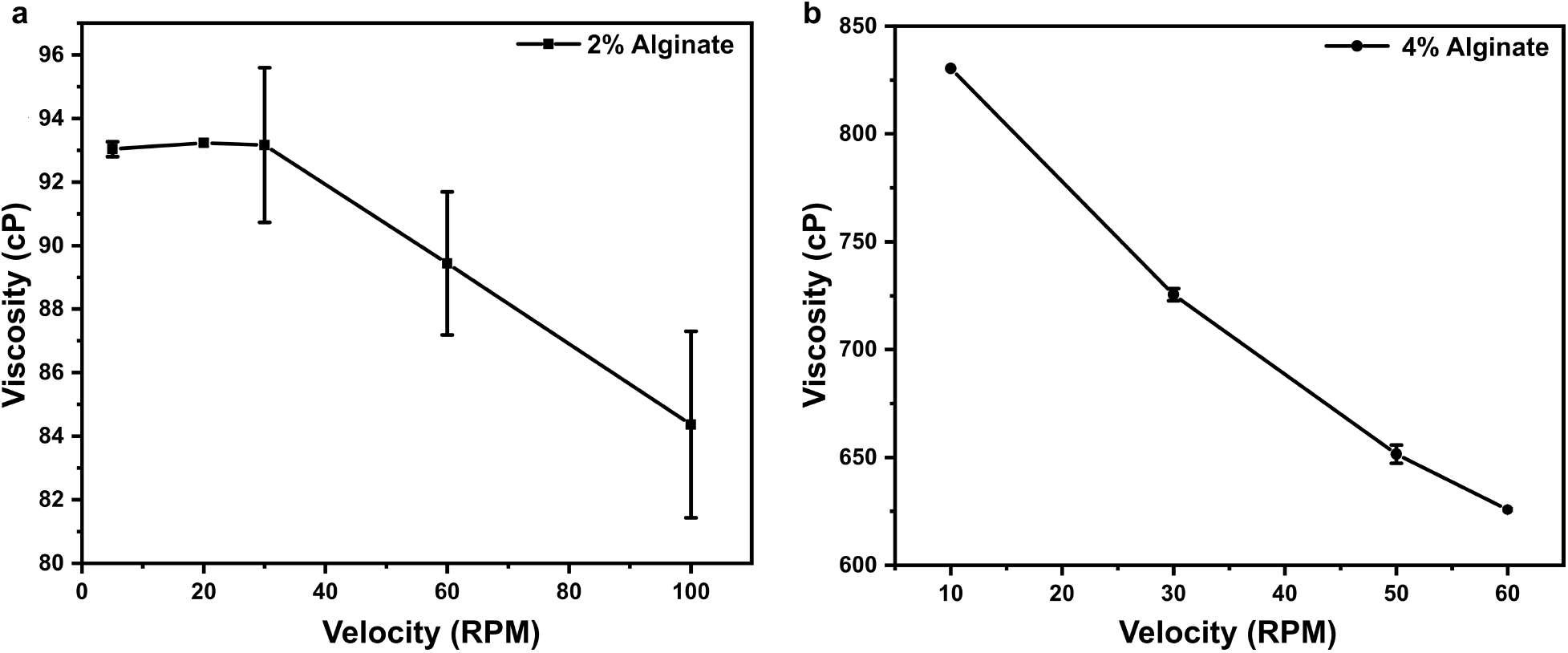
Plots representing viscosities of alginate solution and their variation with velocities. (a) Dynamic viscosity of 2 % sodium alginate solution in DI water at different velocities. (b) Variation of dynamic viscosity of 4 % sodium alginate solution in DI water with velocity. The decreasing trend of the viscosity with an increase in the velocity showed the shear thinning effect exhibited by the alginate solution.

The alginate concentration was increased from 2 % w/v to 4 % w/v while keeping the concentration of the core and sheath solution constant at 15 % w/v PEG and 10 % w/v PEG, respectively, to investigate the displacement of fluorescent particles suspended in the core solution. Four different core flow rates were tested, 100, 150, 200, 250 *μL min*^−1^ to study the effect of flow rates on the particle displacement while keeping the cladding and sheath flow rates constant at 200 and 150 *μL min*^−1^. For this configuration, unlike 2 % w/v alginate, it was observed from the brightfield and corresponding fluorescent images that the outward displacement of the particles was confined at the interface of the core and the cladding solution. **Figure 4 (b)** shows the microparticle seeding density measured using the % seeding area within the brightfield micrographs. The seeding density of the microparticles seeded on the inner surface of the alginate hollow microfibers was observed to increase with the increase in the core flow rate. For flow rate ratio of 250: 200: 150 *μL min*^−1^, the seeding density increased approximately by 192 % as compared to flow rate ratio of 100: 200: 150 *μL min*^−1^. The brightfield and the corresponding fluorescent images of the seeded microparticles at different flow rate ratios depicting the seeded density are shown in **Figure 4 (a)**. The increase in the lateral displacement towards the interface and hence, the seeding density is critical in minimizing the loss of microparticles escaping to the collection bath. This phenomenon is critical in applications where there is an involvement of cells or biological materials in limited quantity.

**Figure 4.**
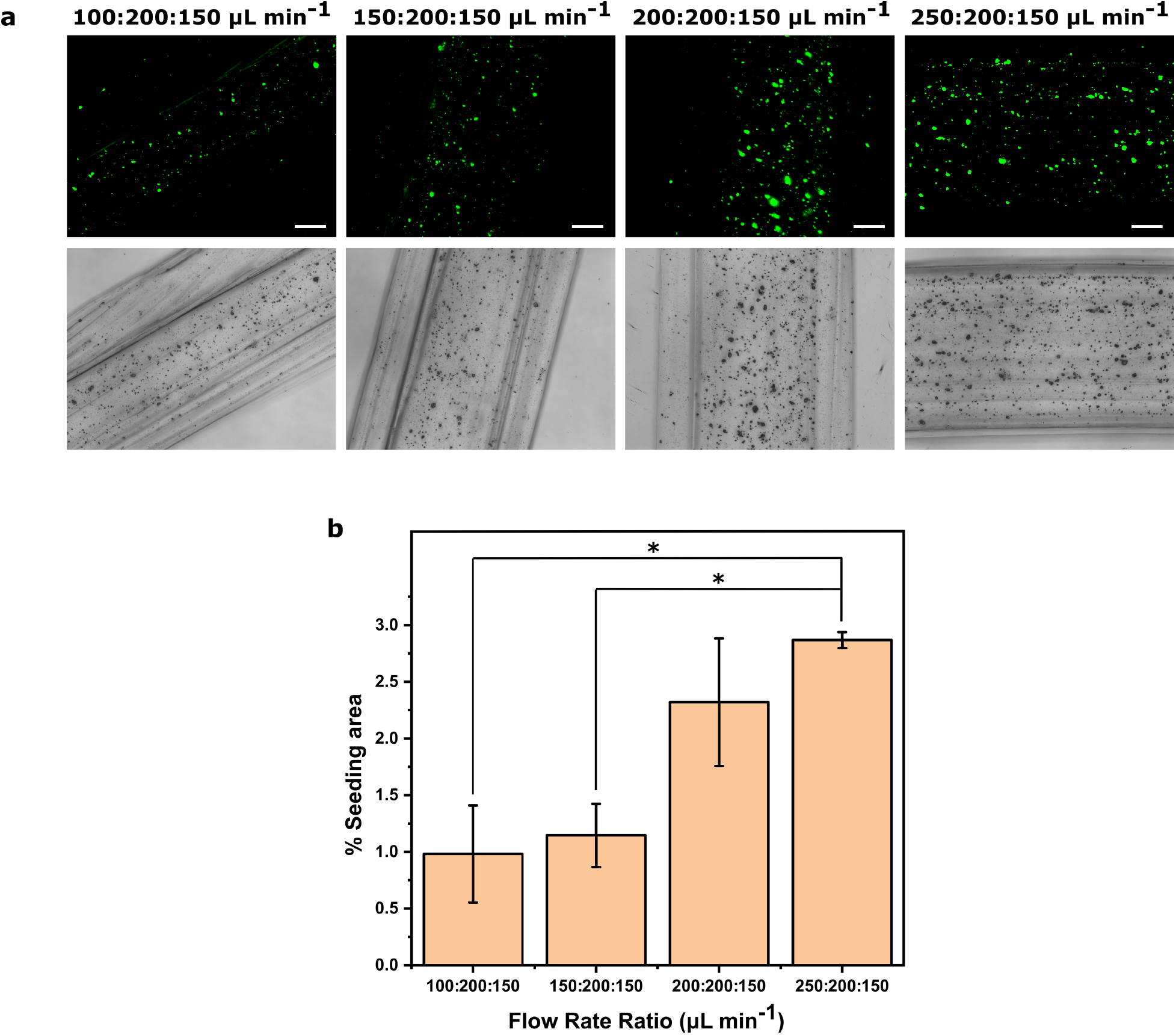
a) Fluorescence and brightfield images of the microparticles seeded on the inner surface of the alginate hollow microfibers. The flow rate ratios used are as follows: 100: 200: 150, 150: 200: 150, 200: 200: 150, 250: 200: 150, all in μL min^−1^. The scale bars are 100 μm. b) Microparticle seeding density measured as % seeding area for the given flow rate ratios. The microparticle seeding density increased with the increase in the core flow rate. ANOVA was performed to test the statistical significance and post-hoc Tukey test determined the significance between specific groups (*p < 0.05).

### Effect of flow rate ratios on the particle displacement and dimensions of the hollow microfibers

Both external and internal widths of the alginate hollow microfibers were investigated using variable flow rate ratios. The flow rate ratios used were 100: 200: 150, 150: 200: 150, 200: 200: 150, and 250: 200: 150 *μL min*^−1^ and the fluorescence images of their corresponding alginate hollow microfibers are shown in **Figure 4 (a)**. The sheath flow rate was set such that the extrusion of the alginate-peg concentric fluid regime and its polymerization with the calcium chloride bath solution did not result in unwanted solidification within the microfluidic channel. Moreover, the cladding flow rate was set in a way that the flow rates for the core investigated could easily displace the former at the hydrodynamic regions within the microchannel architecture. It was observed that the inner width of the alginate hollow microfibers manufactured increased with increasing core flow rates, as shown in **Figure 5 (a)**. The mean inner widths for the above-mentioned flow rate ratios were measured to be 221.21, 342.47, 398.44, and 475.23 μm. Similarly, an increasing trend in the outer width of the alginate hollow microfibers was observed with increasing core flow rate ratios except between 200:200:150 *μL min*^−1^ and 250: 200: 150 *μL min*^−1^, where it decreased, as shown in **Figure 5 (b)**. This decrease in the outer width could be attributed to the core flow rate exceeding a threshold value, after which the amount of cladding solution advected through the chevrons stayed constant. This shows that simply altering the flow rates of the core solution within a particular range furnishes different dimensions of hollow microfibers.

**Figure 5.**
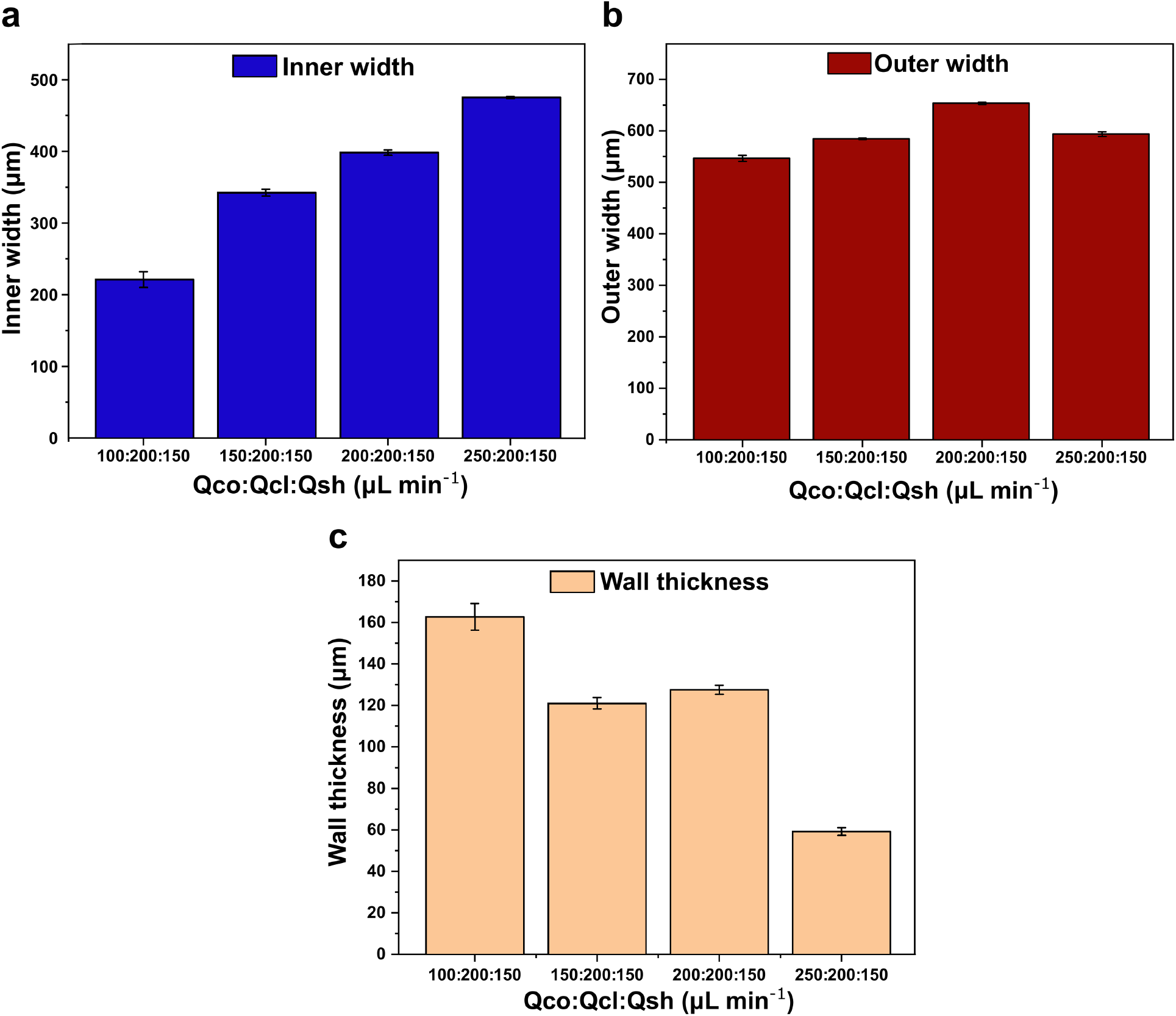
Bar charts representing the (a) inner width, (b) outer width, and (c) wall thickness of the alginate hollow microfibers. The dimensions are plotted with increasing core flow rates while keeping the flow rates of the cladding and sheath solutions constant.

## Conclusions

Microfluidics, along with its many applications, has recently been employed also as an invasive technique to sort or target bio-microparticles flowing through a microchannel for various applications. It utilizes label-free passive forces to direct these microparticles to desired locations within the fluid regime formed in the microfluidic channels. In this study, we use an existing microfluidic cell seeding device used to seed various mammalian cells on the inner surface of polymer-based hollow microfibers to analyze and further enhance the seeding density using fluorescent microparticles resembling as cells. We investigated different concentrations of precursor aqueous sodium alginate and their various flow rate ratios to optimize the overall seeding density. It was observed that the 4 % w/v sodium alginate solution contained the microparticles from displacing across the interface between the core and cladding solutions, unlike its 2 % w/v counterpart, where the microparticles were also observed to be encapsulated in microfiber walls. Moreover, it was observed that by employing higher core flow rate of 250 *μL min*^−1^ increased the microparticle seeding density by 192 % compared to lower flow rate of 100 *μL min*^−1^. This increase in the cell density could be attributed to the higher lateral displacement of microparticles towards the walls at higher core flow rates. The results obtained could be translated to minimize the loss of the cells or bio-microparticles during their seeding on the inner surface of the polymer hollow microfibers, expediting the manufacturing of the in-vitro setups potentially used for drug delivery applications.

## Experimental Section

### Materials

Polyethylene glycol (PEG, Mw = 20,000) (8-18897, Sigma-Aldrich) was used as core and sheath solutions. Sodium Alginate (A2033, Sigma-Aldrich) was used as a precursor solution for the manufacturing of hollow microfibers. Calcium Chloride Dihydrate (CaCl_2_) (C79-3, Thermo Fischer Scientific) solution was used as a collection bath and acted as a polymerizing agent for the solidification of the alginate-based hollow microfibers. Fluorescent microparticles (MMI-105, Bioclone) with 1 *μ*m diameter were used for seeding on the inner surface of the alginate hollow microfibers. Polymethyl methacrylate (PMMA), commonly known as Acrylic, was bought from Grainger, and the 0.2 mm diameter two-flute end mills were from Harvey Tools.

### Microfluidic device fabrication

The detailed procedure of fabricating the microfluidic device used for seeding microparticles on the inner surface of the alginate hollow microfibers is reported in our previous articles. ^32, 36^ Briefly, a unique microchannel architecture was designed to model different polymer solutions into a concentric fluid regime. The design consists of 5 inlet channels, namely, a singular core inlet, a pair of cladding inlets, and a pair of sheath inlets converging on a central channel. Moreover, it comprises of two hydrodynamic focusing regions, where the cladding and the sheath solutions converge on the central channel, and two sets of chevrons, which are v-shaped grooves responsible for inducing advection of the outer layer of the fluid. The microchannel design was milled on a polymethyl methacrylate (PMMA), commonly known as Acrylic, chips using a computer numerical control (CNC) micro mill. Acrylic was chosen as a substrate as it is rigid, meaning it wouldn’t interfere with the flow modulation within the microfluidic device, and it is transparent, which is good for optical microscopy. The chips were plasma treated and subsequently bonded together using a thermally-solvent assisted bonding technique to obtain a microfluidic device ^37^.

### Manufacturing fluorescent microparticles seeded alginate hollow microfibers

2 % w/v or 4 % w/v Sodium Alginate in di-ionized water (DI) was infused through the cladding inlets as a precursor, while 10 % w/v polyethylene glycol (PEG) in DI was infused through the sheath inlets as a template fluid. The fluorescent microparticles with a concentration of 17 × 10^6^ microparticles/ml were suspended in the core solution containing 15 % w/v PEG to be seeded on the inner surface of the alginate hollow microfibers. The seeding density was analyzed and further optimized using different flow rates and concentrations of the core and cladding solutions.

### Imaging

Immediately after manufacturing the fluorescent microparticles seeded alginate hollow microfibers using different concentrations of alginate solution and flow rate ratios, they were imaged using an inverted fluorescence microscope. Z-stack imaging was used to image all the seeded microparticles along the height of the alginate hollow microfibers. The z-stack of the images was stacked together using ImageJ software to investigate the location of the seeded microfibers along their width.

### Viscosity measurement

The viscosity of the alginate solution was measured using a Brookfield DV1 Viscometer. The viscosity of 2 % w/v and 4 % w/v alginate solution was measured at different rotational speeds to analyze the change in the viscosity. The shear thinning effect exhibited by the alginate solution was quantified by calculating a shear thinning index (m), which is given by the apparent viscosity of the fluid at a higher rotational speed divided by the apparent viscosity at a lower rotational speed.

### Statistical Analysis

To test the statistical significance, ANOVA was performed and a significance value of p < 0.05 was considered. Tukey’s test for post-Hoc analysis was used to determine the significant difference between specific groups.

## Acknowledgment

This work was partially supported by the National Science Foundation Award 2014346.

## Data Availability

The data that support the findings of this study are openly available in Mendeley Data at https://data.mendeley.com/drafts/8tfp86wwcx, reference number V2 ^38^.

